# KOunt – A reproducible KEGG orthologue abundance workflow

**DOI:** 10.1101/2023.04.27.538265

**Authors:** Jennifer Mattock, Marina Martínez-Álvaro, Matthew A. Cleveland, Rainer Roehe, Mick Watson

**Affiliations:** The Roslin Institute and Royal (Dick) School of Veterinary Studies, University of Edinburgh, Edinburgh, Midlothian, UK; Scotland’s Rural College, Edinburgh, UK; Genus plc, DeForest, WI, USA

## Abstract

**Summary:** Accurate gene prediction is essential for successful metagenome analysis. We present KOunt, a Snake-make pipeline, that precisely quantifies KEGG orthologue abundance.

**Availability and implementation:** KOunt is available on GitHub: https://github.com/WatsonLab/KOunt. The KOunt reference database is available on figshare: https://figshare.com/s/72fe3ed7dfbbdcd8b1e5. Test data are available at https://figshare.com/ndownloader/files/39545968 and version 1.1.0 of KOunt at https://figshare.com/ndown-loader/files/39547273.

**Supplementary information:** Supplementary data are available at *Bioinformatics* online.

## 1 Introduction

Accurate and effective sequence annotation is key in interpreting metagenomic sequence data. The KEGG database is a popular reference database that groups proteins into functional orthologues, termed KEGG orthologues (KOs) (Kanehisa *et al*., 2021). Several tools that identify KO abundance exist with varying aims. FMAP is a functional analysis pipeline that aligns reads to a KEGG filtered UniProt reference database and calculates gene family abundance (Kim *et al*., 2016). DiTing uses KofamKOALA to identify KOs and calculates relative abundance (Xue *et al*., 2021). Both HumanN2 and Metalaffa provide conversion between UniRef90 hits and KOs; HumanN2 also allows searching against a legacy version of the KEGG database (Franzosa *et al*., 2018; Eng *et al*., 2020).

Here we describe KOunt, a reproducible workflow which uses freely available software to calculate KO abundance in metagenomic sequence data, taking multiple approaches to improve the annotation of proteins and reads that initially do not have a hit. Unlike other KO abundance tools, KOunt gives the user the option to calculate the abundance of the RNA KOs in the metagenomes and also cluster the proteins by sequence identity to report the diversity within each KO. KOunt has been used to successfully quantify KO abundance in rumen microbiome samples (Martínez-Álvaro *et al*., 2022).

## 2 Features

KOunt uses Snakemake to generate a scalable, reproducible workflow, utilising freely available software (Köster and Rahmann, 2012). The pipeline is accompanied by reads subsampled from ERR2027889 to quickly test that installation has completed successfully. Reads are trimmed, assembled, proteins predicted and coverage calculated with Fastp, Megahit, Prodigal and BEDTools respectively (Hyatt *et al*., 2010; Quinlan and Hall, 2010; Li *et al*., 2015; Chen *et al*., 2018). Complete proteins are annotated with a KO using KofamScan (Aramaki *et al*., 2020). These proteins are subsequently clustered by 100%, 90% and 50% sequence identity with CD-Hit and MMseqs2 to quantify the diversity within each KO (Li and Godzik, 2006; Steinegger and Söding, 2017).

Users then have the option of using the custom KOunt databases to further annotate proteins and reads without a hit. Proteins are blasted against the KOunt protein database with Diamond (Buchfink, Reuter and Drost, 2021) and then assessed for RNA presence using Barrnap (https://github.com/tseemann/barrnap/) and tRNAscan-SE 2.0 (Chan *et al*., 2021). All unannotated reads are then screened against the KOunt RNA database with MMseqs2 and kallisto (Bray *et al*., 2016). An indepth description of the pipeline is available in Supplementary Information.

DiTing both missed high abundance KOs and overestimated several, such as K07497 whose abundance increased from 294,342 in the ground truth results to 483,177. FMAP had a better correlation to the ground truth (*r*=0.87 ± 0.002) but was still missing many high abundance KOs. KOunt was able to annotate the high abundance KOs missed by the other approaches. Many of these were RNA, which KOunt accurately quantified unlike DiTing and FMAP. Of the 12,945 KOs present in the reads according to the KEGG annotation, KOunt found the most at 11,466, followed by FMAP with 10,735 and DiTing with 9,681. Whilst KOunt performed the best at identifying KOs reported in the ground truth, it also found the largest number of KOs (1,608) not reported by the ground truth, versus 1,228 and 188 by FMAP and DiTing respectively. This could indicate that KOunt finds more false positives than the other approaches, however we think it’s likely that, due to the multitude of approaches KOunt uses to quantify proteins, KOunt is identifying proteins that haven’t yet been described in the KEGG annotation.

**Figure 1.**
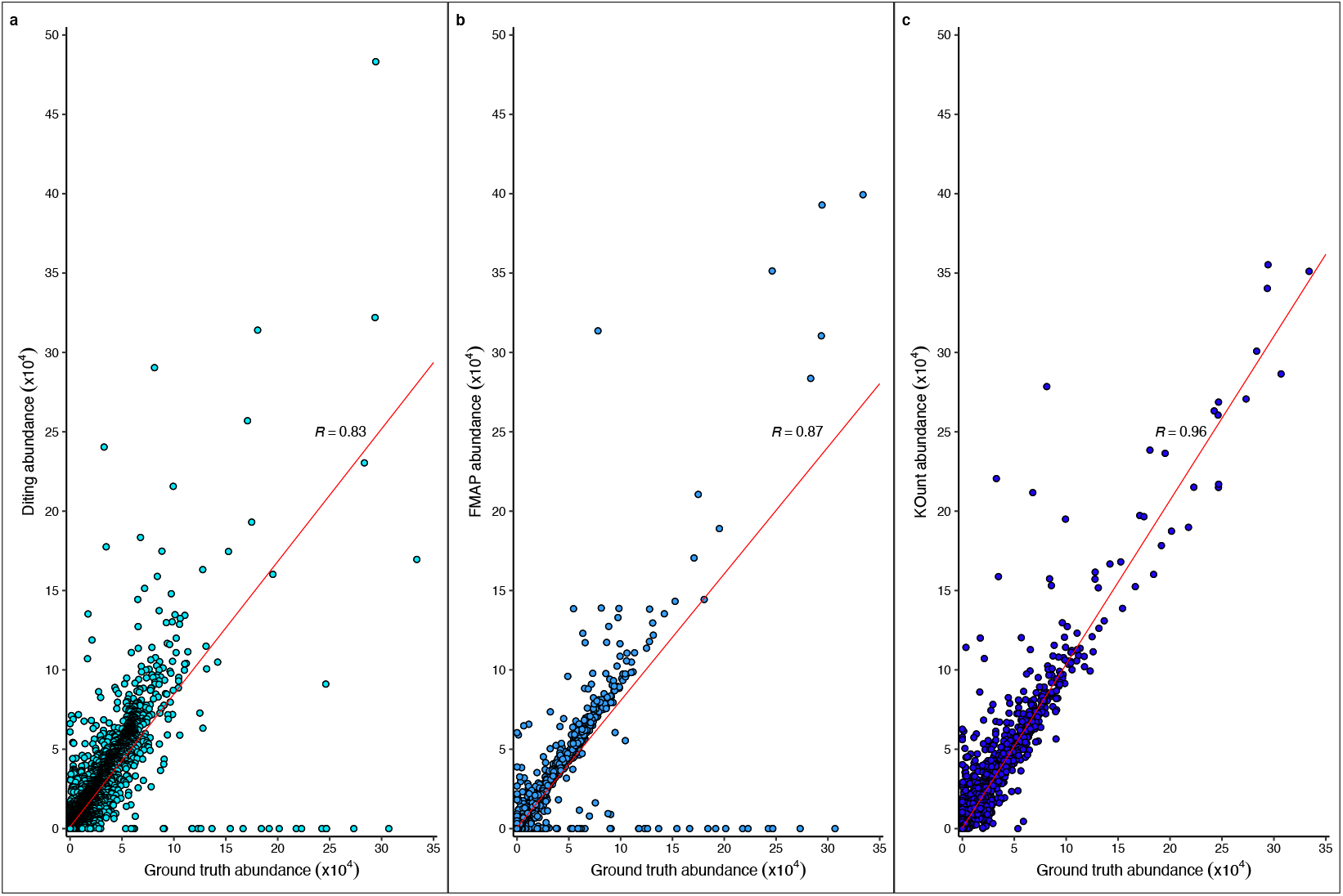
Comparison of ground truth KO abundance and DiTing, FMAP and KOunt KO abundance. a) Ground truth versus DiTing KO abundance b) Ground truth versus FMAP KO abundance c) Ground truth versus KOunt KO abundance

Many proteins from microbiomes are unable to be annotated, for example 40% of the 170 million proteins in the Unified Human Gastrointestinal Genome collection are unannotated (Almeida *et al*., 2021). Therefore, retaining as many reads as possible while maintaining accuracy is paramount. Across the ten samples FMAP and DiTing used on average 78 million and 79 million reads respectively; KOunt outperformed both, capturing an average of 89 million reads per sample. Whilst this is clearly beneficiary, as 150 million reads were inputted there is still a need for improved protein annotation of reference datasets.

KOunt also clusters the proteins identified by KofamScan by sequence identity, allowing investigation of the diversity within KOs. In this dataset three million proteins were identified, which grouped into 0.4 million 90% clusters and 0.2 million 50% clusters. K03406, methyl-accepting chemotaxis proteins, was the KO with the largest number of 50% clusters (1311) identified with KOunt; as a protein needs to have just 50% similarity to one of the proteins in a cluster to be included in that cluster, this illustrates the vast amount of diversity within this KO. The grouping of homologous proteins enables further investigation of highly abundant clusters and those with abundance associated with traits of interest.

To conclude we present KOunt, a reproducible, scalable pipeline which accurately calculates raw KO abundance from metagenomic sequencing reads. Furthermore, KOunt also reports the number of 90% and 50% sequence identity clusters in each KO, showing the protein diversity within the KOs and facilitating exploration of groups of unannotated proteins.

## Acknowledgements

This work has made use of the resources provided by the Edinburgh Compute and Data Facility (ECDF) (http://www.ecdf.ed.ac.uk/).

## Data availability

The KOunt pipeline is available at https://github.com/WatsonLab/KOunt. The KOunt reference database is available on figshare: https://figshare.com/s/72fe3ed7dfbbdcd8b1e5. Test data are available at https://figshare.com/ndownloader/files/39545968 and version 1.1.0 of KOunt at https://figshare.com/ndownloader/files/39547273.

