## Supplementary Information for "KOunt – A reproducible KEGG orthologue abundance workflow"

**Workflow description**

Firstly, raw sequencing reads are trimmed with fastp to remove adaptors, reads below a minimum length cut-off and polyG tails (Chen *et al.*, 2018). The trimmed reads ids are shortened with FastUtils (https://github.com/nunofonseca/fastq_utils) and the reads assembled using MEGAHIT (Li *et al.*, 2015), proteins predicted using Prodigal (Hyatt *et al.*, 2010) and filtered to remove incomplete proteins. To calculate protein abundance BWA-MEM is used to map the trimmed reads to their assembly (Li, 2013). Coverage computation speed is increased using BamDeal (https://github.com/BGI-shenzhen/BamDeal) to split the BAM file into groups, BEDTools is then utilised to calculate protein coverage for each split BAM file before the outputs for each sample are combined (Quinlan and Hall, 2010).

KOunt uses KofamScan to annotate the complete proteins of each sample (Aramaki *et al.*, 2020). Hits that don’t pass KofamScan’s thresholds for each KO are discarded and are labelled ‘NoHit’. Proteins that are annotated with more than one KO have the option of having their coverage split between these KOs or classified as ‘Multiple’, enabling further analysis of these proteins. The coverage results are combined with the KofamScan results and the abundance of each KO, NoHit and Multiple in each sample calculated; the results for each sample are combined to produce a KO abundance matrix. Proteins from each KO are clustered at 100% sequence identity with Cd-hit (Li and Godzik, 2006), and then at 90% and 50% sequence identity with MMseqs2 (Steinegger and Söding, 2018). To quantify the diversity within each KO, Multiple and NoHit, the number of clusters are counted and outputted.

Users then have the option to annotate unannotated proteins and reads with a custom KO reference database. As described in Kim *et al*., 2016, UniProt proteins from archaea, bacteria and fungi, that had been annotated with a KO, were downloaded in January 2022 (Kim *et al.*, 2016). Additionally, proteins in the UniProt cross-referenced database (<https://www.uniprot.org/database?query=*>) that had a KEGG gene id and had been reviewed were filtered to include only prokaryotic and eukaryotic microorganisms. The R package KEGGREST was used to convert the proteins with EC ids into KOs initially and then the remaining KEGG gene ids into KOs (Tenenbaum, 2020). Proteins that belonged to multiple KOs were annotated with all of these KOs. To create an RNA KO database the following reference sequences were downloaded: the small and large subunit 99% dereplicated and truncated SILVA reference database (v138.1)(Quast *et al.*, 2013); the GtRNA database (downloaded 04/22)(Chan and Lowe, 2009); all RNA in the KEGG genes database (downloaded 06/22)(Kanehisa *et al.*, 2016) and the RNA families in Rfam (downloaded 04/22)(Kalvari *et al.*, 2021). The tRNA family RF00005 was processed with tRNAscan-SE 2.0 (Chan *et al.*, 2021) to identify the type of each tRNA.

NoHit proteins are blasted against the KOunt protein database with Diamond and the RNA database with MMseqs2 (Steinegger and Söding, 2017; Buchfink, Reuter and Drost, 2021). Those with a percentage identity greater than 80% and a query coverage 90% <> 110% are identified as the KO of their top hit; if a protein has an identical percentage identity and e-value to multiple KOs then it is annotated with all of these KOs. If RNA abundance is desired the remaining Nohit proteins are processed by Barrnap (<https://github.com/tseemann/barrnap/>) and tRNAscan-SE 2.0; again if a protein has multiple hits KOunt annotates it with all.

KOunt then allows the user to annotate unmapped reads, those that mapped to incomplete proteins and those mapped to NoHit proteins with the KOunt databases. The reads are blasted against the protein database using Diamond and the RNA database with MMseqs2 (Steinegger and Söding, 2017; Buchfink, Reuter and Drost, 2021). The results are combined, hits with a percentage identity of less than 80% are removed and the top hit, when sorted by e-value and percentage identity, is determined for each read. If a read has multiple top hits with the same e-value and percentage identity, then they are all taken forward and coverage is split between them, preventing biased mapping to KOs occurring. Mean coverage is then calculated using the coordinates of the read, length of the reference protein and the number of top hits. All reads that remain unannotated are processed with kallisto and the KOunt RNA database to quantify RNA abundance (Bray *et al.*, 2016).

**Benchmarking methods**

To generate ground truth data, genomes from the Hungate1000 Collection and the Human Gastrointestinal Bacteria Culture Collection (PRJNA482748) were compared to the KEGG organisms database (downloaded in 2019) using MASH (Ondov *et al.*, 2016; Seshadri *et al.*, 2018; Zou *et al.*, 2019). The KEGG genomes with the smallest mash distance to each of the microbiome genomes were deduplicated, resulting in a collection of 323 genomes. Metagenomic reads for these genomes were simulated ten times, using InSilicoSeq with the options draft, basic, n_reads 150000000 and varying seed between 1-10 for each repeat (Gourlé *et al.*, 2019). KEGGREST was used to download coordinates for the KOs in the KEGG genomes (Tenenbaum, 2020). Ground truth KO abundance was calculated using BEDTools (version 2.30.0), with the coverage of areas with multiple hits split between the KOs and the coverage of genes that were reported as split into several locations calculated as the mean read depth of all the locations (Quinlan and Hall, 2010).

These simulated reads were used as input for KOunt, without the default options, and trimmed with fastp for use with DiTing and FMAP, both using default parameters (Kim *et al.*, 2016; Chen *et al.*, 2018; Xue *et al.*, 2021). As the desired output was KO abundance the FMAP meanDepth results were used. As DiTing produces normalised KO abundance the raw abundance was manually calculated using the BBMap pileup and KEGG annotation files.

The Pearson correlation coefficients between the ground truth KO abundance and those produced by KOunt, DiTing and FMAP were calculated using R (version 4.1.0) and the plot generated with ggplot2 (version 3.3.6), ggpubr (version 0.5) and cowplot (version 1.1)(Wickham, 2009; R Core Team, 2018; Wilke, 2020; Kassambara, 2022).

KOunt is accompanied by some FASTQs to test that installation of the conda environments has completed successfully. These reads were subsampled from a rumen microbiome metagenome, ERR2027889, using seqtk (<https://github.com/lh3/seqtk>) with seed of 1234 and proportion of 0.005.
